# Chromosome-scale assembly comparison of the Korean Reference Genome KOREF from PromethION and PacBio with Hi-C mapping information

**DOI:** 10.1101/674804

**Authors:** Hui-Su Kim, Sungwon Jeon, Changjae Kim, Yeon Kyung Kim, Yun Sung Cho, Jungeun Kim, Asta Blazyte, Andrea Manica, Semin Lee, Jong Bhak

## Abstract

**Background:** Long DNA reads produced by single molecule and pore-based sequencers are more suitable for assembly and structural variation discovery than short read DNA fragments. For *de novo* assembly, PacBio and Oxford Nanopore Technologies (ONT) are favorite options. However, PacBio’s SMRT sequencing is expensive for a full human genome assembly and costs over 40,000 USD for 30x coverage as of 2019. ONT PromethION sequencing, on the other hand, is one-twelfth the price of PacBio for the same coverage. This study aimed to compare the cost-effectiveness of ONT PromethION and PacBio’s SMRT sequencing in relation to the quality.

**Findings:** We performed whole genome *de novo* assemblies and comparison to construct an improved version of KOREF, the Korean reference genome, using sequencing data produced by PromethION and PacBio. With PromethION, an assembly using sequenced reads with 64x coverage (193 Gb, 3 flowcell sequencing) resulted in 3,725 contigs with N50s of 16.7 Mbp and a total genome length of 2.8 Gbp. It was comparable to a KOREF assembly constructed using PacBio at 62x coverage (188 Gbp, 2,695 contigs and N50s of 17.9 Mbp). When we applied Hi-C-derived long-range mapping data, an even higher quality assembly for the 64x coverage was achieved, resulting in 3,179 scaffolds with an N50 of 56.4 Mbp.

**Conclusion:** The pore-based PromethION approach provides a good quality chromosome-scale human genome assembly at a low cost with long maximum contig and scaffold lengths and is more cost-effective than PacBio at comparable quality measurements.

## Data Description

Next-generation sequencing (NGS) is a set of powerful sequencing technologies and a recent trend in genomics is to use cost-effective long DNA reads for assembly and structural variation discovery using single molecule sequencing methods. Oxford Nanopore Technologies (ONT) and PacBio platforms have advantages of a short run time and long read lengths over short fragmented reads by Illumina [1, 2]. Unfortunately, both methods share high base-calling error rates [3, 4]. However, bioinformatics pipelines for self-error correction and/or polishing sequences with short reads have become an effective option, and the overall accuracy of long read based assemblies is approaching what is required to be a viable option for personal reference genome construction [5]. Despite its excellent performance, PacBio’s SMRT sequencing is expensive for the effective coverage required for a full human genome assembly, costing over 40,000 USD for 30x coverage (with 15 SMRT cells; from an estimated 6 Gbp raw reads production per SMRT cell) as of 2019 [6, 7, 8]. On the other hand, the nanopore based single molecule, long read platform, PromethION from ONT is highly cost-effective at one-twelfth the price of PacBio’s for the same read amount, with an advantage of even longer average and maximum read lengths [9]. Although the two methods share some similarity, they are fundamentally different in that ONT uses a minimal amount of reagents with small form factor devices, and can be a promising future technology for a very broad scope of applications given its advantageous size and cost.

In this study, we performed benchmark tests of PromethION and PacBio with low and high coverages of sequencing data and investigated the advantages of pairing these long read technologies with very long-range chromosome mapping information by Hi-C, using the already existing high-quality Korean reference genome, KOREF, as a benchmark [10].

## Whole genome sequencing by ONT PromethION R9.4.1 platform

Human KOREF cell lines (http://koref.net) were cultured at 37°C in 5% CO_2_ in RPMI-1640 medium with 10% heat-inactivated fetal bovine serum. DNA was extracted from cells using the DNeasy Blood & Tissue kit (Qiagen). The KOREF cells (5 × 10^6^) were centrifuged at 300 *g* for 5 min; the pelleted cells were suspended in 200 μL of PBS and DNA was extracted according to the manufacturer's instructions. To preserve large-sized DNA and purify DNA fragments, we used Genomic DNA Clean & Concentrator kit (Zymo). The DNA quality and size were assessed by running 1 μL of purified DNA on the Bioanalyzer system (Agilent). Concentration of DNA was assessed using the dsDNA BR assay on a Qubit fluorometer (Thermo Fisher).

DNA repair (NEBNext FFPE DNA Repair Mix, NEB M6630) and end-prep (NEBNext End Repair/dA-tailing, NEB E7546) were performed using 1 μg human genomic DNA. The mixture of 1 μL DNA CS, 3.5 μL FFPE Repair Buffer, 2 μL FFPE DNA Repair Mix, 3.5 μL Ultra II End-prep reaction buffer, and 3 μL Ultra II End-prep enzyme mix was added to 47 μL DNA sample. The final mixture was incubated at 20°C for 5 min and then at 65°C for 5 min, cleaned up using 60 μL AMPure XP beads, incubated on Hula mixer for 5 min at room temperature, and washed twice with 200 μL fresh 70% ethanol. The pellet was allowed to dry for 30 s, and then DNA was eluted in 61 μL of nuclease-free water. An aliquot of 1 μL was quantified by Qubit to ensure ≥ 1 μg DNA was retained.

Adaptor ligation was performed by adding 5 μL of Adaptor Mix (AMX, SQK-LSK109 Ligation Sequencing Kit 1D, Oxford Nanopore Technologies (ONT)), 25 μL Ligation Buffer (LNB, SQK-LSK109), and 10 μL NEBNext Quick T4 DNA Ligase (NEB, E6056) to 60 μL bead cleaned-up DNA, followed by gentle mixing and incubation for 10 min at room temperature.

The adaptor-ligated DNA was cleaned up by adding 40 μL of AMPure XP beads, incubating for 5 min at room temperature and re-suspending the pellet twice in 250 μL L Fragment Buffer (LFB, SQK-LSK109). The purified ligated DNA was re-suspended in 25 μL of Elution Buffer (ELB, SQK-LSK109), incubated for 10 min at room temperature, followed by pelleting the beads, and transferring the supernatant (pre-sequencing mix or PSM) to a new Eppendorf Lobind tube. A 1-μL aliquot was quantified by Qubit to ensure ≥ 500 ng DNA was retained.

To load the library, 75 μL of Sequencing Buffer (SQB, SQK-LSK109) was mixed with 51 μL of Loading Beads (LB, SQK-LSK109) and this mixture was added to 24 μL DNA library. This library was mixed by pipetting slowly and 150 μL of sample was loaded through the inlet port.

## Whole genome sequencing by PacBio Sequel platform

Genomic DNA was extracted from human KOREF blood samples using QIAGEN Blood & Cell Culture DNA Kit (cat no 13323). A total of 5 μg of each sample was used as input for library preparation. The SMRTbell library was constructed using SMRTbell® Express Template Preparation Kit (101-357-000). Using the BluePippin Size selection system we removed the small fragments for large-insert library. After sequencing primer v4 was annealed to the SMRTbell template, DNA polymerase was bound to the complex (Sequel Binding kit 2.0). We purified the complex using AMPure Purification to remove excess primer and polymerase prior to sequencing. The SMRTbell library was sequenced using SMRT cells (Pacific Biosciences) using Sequel Sequencing Kit v2.1 and 10 h movies were captured for each SMRT Cell 1M v2 using the Sequel (Pacific Biosciences) sequencing platform.

## Short read sequencing by Illumina HiSeq

Short paired-end raw reads using Illumina HiSeq 2000 platform were acquired from a previous study, accession no. SRR2204706 (ftp://ftp.sra.ebi.ac.uk/vol1/srr/SRR220/006/SRR2204706).

## Hi-C chromosome conformation captured reads sequencing

Long distance Hi-C chromosome conformation capture data were generated using the Arima-HiC kit (A160105 v01), and double restriction enzymes were used for chromatin digestion. To prepare KOREF cell line samples for Hi-C analysis, cells were harvested and cross-linked as instructed by the manufacturer. One million cross-linked cells were used as input in the Hi-C protocol. Briefly, chromatin from cross-linked cells or nuclei was solubilized, and then digested using restriction enzymes A1 and A2. The digested ends were then labeled using a biotinylated nucleotide, and ends were ligated to create ligation products. Ligation products were purified, fragmented, and selected by size using AMpure XP Beads. Biotinylated fragments were then enriched using Enrichment beads, and Illumina-compatible sequencing libraries were constructed on End Repair, dA-tailing, and Adaptor Ligation using a modified workflow of the Hyper Prep kit (KAPA Biosystems, Inc.). The bead-bound library was then amplified, and amplicons were purified using AMpure XP beads and subjected to deep sequencing.

## Short and long sequence reads processing

A total of 144 Gbp of short paired-end DNA raw reads were obtained from SRA2204706. Adapter sequences were trimmed from sequenced raw reads using Trimmomatic v0.36 [11] (ILLUMINACLIP:2:30:10 LEADING:5 TRAILING:5 SLIDINGWINDOW:4:20 HEADCROP:15 MINLEN:60) (Trimmomatic, RRID:SCR_011848), and screening for vectors and microbial contaminants were performed using customized database from Refseq. After preprocessing, a total of 137 Gbp cleaned reads were obtained.

A total of 80.7 Gbp and 193 Gbp raw reads (27x and 64x coverage) were obtained as a result of PromethION nanopore sequencing using one and three flowcells. Removing adapter sequences from the raw reads was performed using Porechop v0.2.4 (Porechop, RRID:SCR_016967) [12]. [12]. We also acquired 92.2 Gbp and 187.9 Gbp raw reads from PacBio Sequel sequencing resulting in 30x and 62x coverage (Table 1).

**Table 1.**
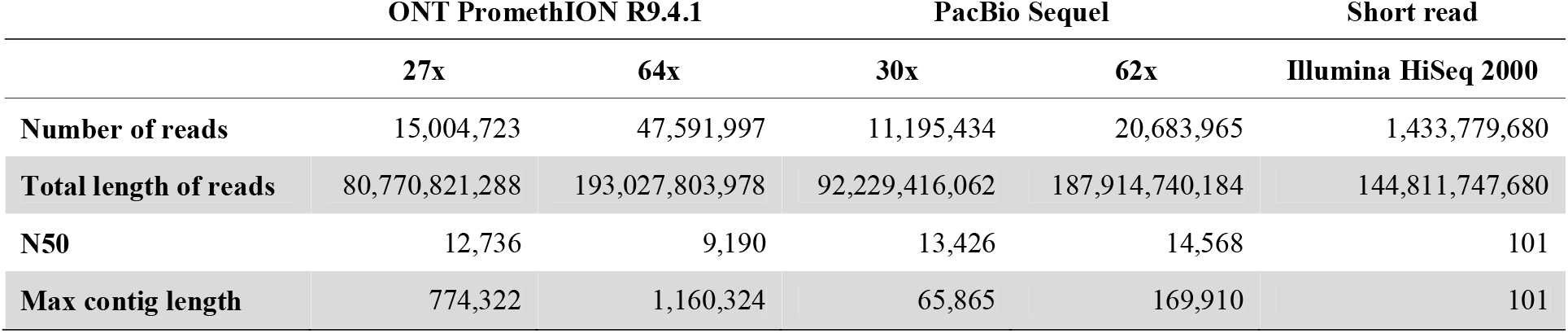
Statistics of raw sequenced reads.

|  | ONT PromethION R9.4.1 |  | PacBio Sequel |  | Short read |
| --- | --- | --- | --- | --- | --- |
|  | 27x | 64x | 30x | 62x | Illumina HiSeq 2000 |
| Number of reads | 15,004,723 | 47,591,997 | 11,195,434 | 20,683,965 | 1,433,779,680 |
| Total length of reads | 80,770,821,288 | 193,027,803,978 | 92,229,416,062 | 187,914,740,184 | 144,811,747,680 |
| N50 | 12,736 | 9,190 | 13,426 | 14,568 | 101 |
| Max contig length | 774,322 | 1,160,324 | 65,865 | 169,910 | 101 |

## Long-read sequence based *de novo* genome assemblies

*De novo* assemblies for the 27x and 64x PromethION raw reads were performed using wtdbg2 v2.3 (WTDBG, RRID:SCR_017225) [13]. To compare the accuracy, two sets of raw reads with 30x and 62x coverage of PacBio Sequel were also used employing the same assembler. Parameters for the assembler were set optimally for each sequencing platform with multiple trials (https://github.com/macarima/KOREF_PromethION_paper). For self-error correction with long reads, we generated consensus sequences using Racon v1.3.2 [14]. To improve the accuracy of assemblies, polishing consensus sequences with 48.2x coverage short reads was performed using Pilon v1.23 v1.23 (Pilon, RRID:SCR_014731) [15]. To assess the completeness of the long-read genome assemblies, BUSCO v3.0.2 (BUSCO, RRID:SCR_015008) [16] with the default AUGUSTUS model for human was used to locate the presence and absence of 4,104 single copy orthologous genes from mammalian OrthoDB v9.

For constructing chromosome-scale assemblies for the PromethION long-reads data, map assembly with Hi-C reads was performed using SALSA2 v2.2 [17]. Duplicated Hi-C reads were removed using clumpify.sh program from BBTools suite v38.32 (Bestus Bioinformaticus Tools, RRID:SCR_016968) [18]. Mapping Hi-C reads to the assembled genome was conducted using the pipeline provided by Arima-Genomics (https://github.com/ArimaGenomics/mapping_pipeline).

Long read assemblies from 27x and 64x PromethION sequencing yielded total assembly sizes of 2,757 Mbp and 2,827 Mbp, with scaffold N50s of 7.6 Mbp and 16.7 Mbp, respectively (Table 2). Assemblies from PacBio sequencing at 30x and 62x coverage yielded the total assembly sizes of 2,800 Mbp and 2,815 Mbp, with scaffold N50s of 11.1 Mbp and 17.9 Mbp, respectively. Adding Hi-C reads to assemblies led to 3.4- to 4.3-fold increase in the scaffold N50 lengths of PromethION (32.7 Mbp for 27x coverage and 56.4 Mbp for 64x coverage). For the PacBio assemblies, 2.2- to 3.3-fold increase was achieved for the scaffold N50 lengths (38.1 Mbp for 30x coverage and 59.3 Mbp for 62x coverage). The longest scaffold from both PromethION and PacBio assemblies with Hi-C was two times the length of the assemblies without Hi-C.

**Table 2.** Statistics of KOREF genome assemblies using ONT PromethION and PacBio Sequel sequencing

| ONT PromethION R9.4.1 |  |  |  |  | PacBio Sequel |  |  |  |
| --- | --- | --- | --- | --- | --- | --- | --- | --- |
|  | 27x assembly * | 64x assembly ** | 27x assembly with Hi-C | 64x assembly with Hi-C | 30x assembly | 62x assembly | 30x assembly with Hi-C | 62x assembly with Hi-C |
| Contigs / Scaffolds No. | 3,262 | 3,725 | 2,313 | 3,179 | 2,443 | 2,695 | 1,476 | 2,139 |
| Total length | 2,757,297,803 | 2,827,624,042 | 2,757,776,303 | 2,827,900,542 | 2,800,962,512 | 2,815,311,932 | 2,801,450,512 | 2,815,594,432 |
| Scaffold N50 | 7,655,153 | 16,706,773 | 32,758,624 | 56,457,651 | 11,137,362 | 17,931,968 | 38,113,117 | 59,361,327 |
| Max contig / scaffold length | 60,569,695 | 88,903,341 | 120,666,262 | 175,227,974 | 50,101,007 | 77,816,513 | 126,818,544 | 174,360,016 |
| Gap | 0.00% | 0.00% | 0.02% | 0.01% | 0.00% | 0.00% | 0.02% | 0.01% |
| GC content | 40.82% | 40.81% | 40.82% | 40.81% | 40.90% | 40.92% | 40.90% | 40.90% |

## Comparison between PromethION and PacBio assemblies

The comparison between PromethION and PacBio assemblies without Hi-C mapping information using sequenced reads at 64x coverage showed comparable quality. In terms of N50, the PromethION assembly at 64x coverage yielded 1.5-fold and 0.93-fold longer N50s compared with the PacBio assemblies at 30x and 62x coverage, respectively (Figure 1a). When we compared the longest contigs, the PromethION assembly at 64x coverage yielded 1.7-fold and 1.1-fold length increase compared with the PacBio assemblies at 30x and 62x coverage, respectively (Figure 1b). Comparing the number of scaffolds, PacBio assembly at 30x coverage showed the fewest (2,443) compared with that of PromethION assembly at 64x coverage (3,725).

**Figure 1.**
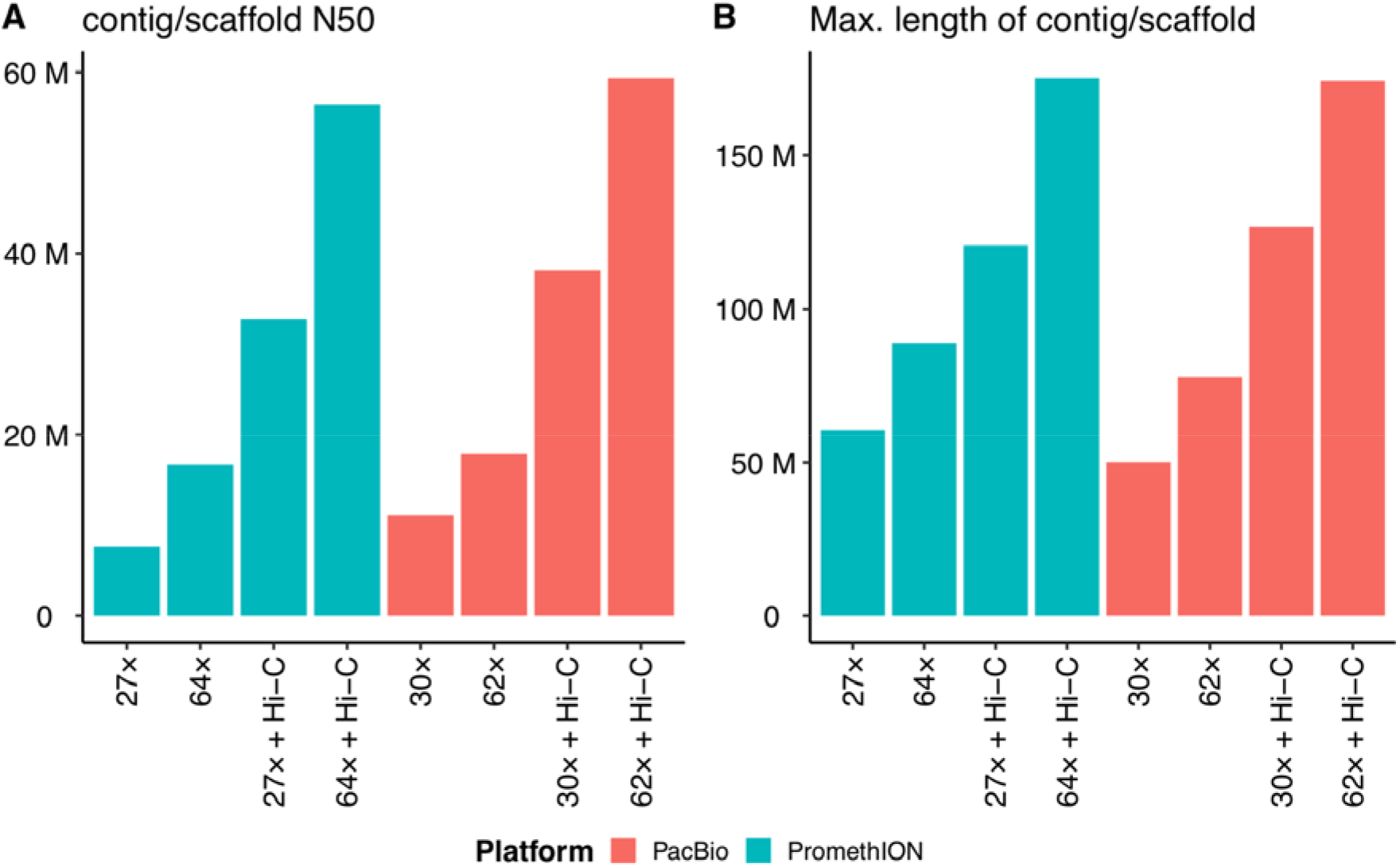
Comparison of N50s and the longest contig/scaffold lengths for PromethION and PacBio assemblies of KOREF

When Hi-C mapping information was added to the assembly construction, the PromethION assembly at 64x coverage showed the best statistics as N50s of 56.4 Mbp and the longest scaffold length of 175.2 Mbp. The PromethION assembly at 27x coverage with Hi-C mapping information yielded 32.7 Mbp for N50s, which was comparable to both 30x and 62x coverage PacBio assemblies with Hi-C; 0.85-fold and 0.55-fold for N50s, respectively (Table 2).

When we compared assessment results from BUSCO, all the assemblies that had been polished with short reads showed good quality; around 92% completed orthologous genes with less than 1.1% completed and duplicated orthologous genes. Comparing the accuracy of the assemblies to the single assembly of KOREF (KOREF_S), which is the current standard, both showed around 99.8% accuracy (Table 3). The accuracy comparison was performed using assess_assembly program from Pomoxis [19].

**Table 3.** Statistics of KOREF genome assembly assessment using BUSCO and accuracy comparison

| BUSCO<br>assessment | ONT PromethION R9.4.1 |  |  |  | PacBio Sequel |  |  |  |
| --- | --- | --- | --- | --- | --- | --- | --- | --- |
|  | 27x assembly | 64x assembly | 27x assembly<br>with Hi-C | 64x assembly<br>with Hi-C | 30x assembly | 62x assembly | 30x assembly<br>with Hi-C | 62x assembly<br>with Hi-C |
| **Complete** | 92.5% | 92.7% | 92.6% | 94.0% | 93.8% | 93.9% | 93.8% | 93.5% |
| **Complete and**<br>**single-copy** | 91.8% | 91.6% | 91.9% | 93.2% | 93.0% | 93.1% | 93.0% | 92.7% |
| **Complete and**<br>**duplicated** | 0.7% | 1.1% | 0.7% | 0.8% | 0.8% | 0.8% | 0.8% | 0.8% |
| **Fragmented** | 3.1% | 3.7% | 3.2% | 3.1% | 3.0% | 2.9% | 3.0% | 3.1% |
| **Missing** | 4.4% | 3.6% | 4.2% | 2.9% | 3.2% | 3.2% | 3.2% | 3.4% |
| **Accuracy**<br>**comparison\*** | 99.78% | 99.73% | 99.78% | 99.73% | 99.83% | 99.79% | 99.86% | 99.80% |

\* Compared with KOREF\_S, the single assembly of KOREF

## Conclusions

We generated high-quality assemblies of the Korean reference genome, KOREF, using ONT’s PromethION long-reads accompanied with Hi-C mapping information and compared them against PacBio sequencing and assemblies of the same sample. Comparing the results from the PromethION 64x sequencing to the PacBio 62x sequencing, we found that the former provided high contiguity and completeness at one-twelfth the cost of PacBio. Results from just 27x PromethION sequencing combined with Hi-C mapping information were also comparable to the 30x and even 62x coverage PacBio sequencing data. Therefore, to generate a chromosome-scale assembly with a long-read technology, at present, the ONT’s PromethION sequencing is a good alternative to PacBio’s, owing to its quality and cost-effectiveness. Simple pore-based long read sequencing has potential to dramatically improve sequencing and subsequent bioinformatics analysis for personal genome projects and cancer genome analyses where *de novo* assemblies are necessary for structural and copy number variations that cannot be detected easily by conventional short read only methods.

## Availability of supporting data

Raw long-read sequencing data from PromethION and PacBio is available at NCBI genbank under the project accession number PRJNA549351. All genome assemblies of KOREF are available at KOREF website (http://koref.net).

## Abbreviations

BUSCO: Benchmarking Universal Single-Copy Orthologs
PacBio: Pacific Biosciences
SMRT: single-molecule real-time

## References

1. Mccarthy A. Third generation DNA sequencing: Pacific biosciences’ single molecule real time technology. Chem Biol [Internet]. Elsevier Ltd; 2010;17:675–6.

2. Laver T, Harrison J, O’Neill PA, Moore K, Farbos A, Paszkiewicz K, et al. Assessing the performance of the Oxford Nanopore Technologies MinION. Biomol Detect Quantif [Internet]. Elsevier GmbH; 2015;3:1–8.

3. Chin C-S, Alexander DH, Marks P, Klammer AA, Drake J, Heiner C, et al. Nonhybrid, finished microbial genome assemblies from long-read SMRT sequencing data. Nat Methods [Internet]. 2013;10:563–9.

4. Laver T, Harrison J, O’Neill PA, Moore K, Farbos A, Paszkiewicz K, et al. Assessing the performance of the Oxford Nanopore Technologies MinION. Biomol Detect Quantif [Internet]. Elsevier GmbH; 2015;3:1–8.

5. Fu S, Wang A, Au KF. A comparative evaluation of hybrid error correction methods for error-prone long reads. Genome Biol [Internet]. Genome Biology; 2019;20:1–17.

6. Desk R. Review article. Toenail onychomycosis: an important global disease burden. [Internet]. N. Engl. J. Med. 2005. p. 1–4.

7. University of Washington PacBio Sequencing Services. Available from: https://pacbio.gs.washington.edu/

8. UC Davis Genome Center. Available from: https://dnatech.genomecenter.ucdavis.edu/prices/

9. Nanopore tech. Available from: https://nanoporetech.com/products/comparison/

10. Cho YS, Kim H, Kim HM, Jho S, Jun JH, Lee YJ, et al. Corrigendum: An ethnically relevant consensus Korean reference genome is a step towards personal reference genomes. Nat Commun [Internet]. 2017;8:16168.

11. Bolger AM, Lohse M, Usadel B. Trimmomatic: A flexible trimmer for Illumina sequence data. Bioinformatics [Internet]. 2014;30:2114–20

12. Porechop, adapter trimmer for Oxford Nanopore reads. https://github.com/rrwick/Porechop

13. Ruan J, Li H. Fast and accurate long-read assembly with wtdbg2. bioRxiv [Internet]. 2019;530972.

14. Racon, ultrafast consensus module for raw de novo genome assembly of long uncorrected reads. https://github.com/isovic/racon

15. Walker BJ, Abeel T, Shea T, Priest M, Abouelliel A, Sakthikumar S, et al. Pilon: An integrated tool for comprehensive microbial variant detection and genome assembly improvement. PLoS One [Internet]. 2014;9.

16. Simão FA, Waterhouse RM, Ioannidis P, Kriventseva E V., Zdobnov EM. BUSCO: Assessing genome assembly and annotation completeness with single-copy orthologs. Bioinformatics [Internet]. 2015;31:3210–2.

17. Ghurye J, Pop M, Koren S, Bickhart D, Chin CS. Scaffolding of long read assemblies using long range contact information. BMC Genomics [Internet]. BMC Genomics; 2017;18:1–11.

18. BBMap, short read aligner and other bioinformatics tools. https://sourceforge.net/projects/bbmap/

19. Pomoxis, bioinformatics tools for nanopore research. https://nanoporetech.github.io/pomoxis/

